# Predicting drug resistance evolution: insights from antimicrobial peptides and antibiotics

**DOI:** 10.1101/138107

**Authors:** Guozhi Yu, Desiree Y Baeder, Roland R Regoes, Jens Rolff

**Author notes:** shared first authors (Guozhi Yu, Desiree Y Baeder).

## Abstract

Antibiotic resistance constitutes one of the most pressing public health concerns. Antimicrobial peptides of multicellular organisms are considered part of a solution to this problem, and AMPs produced by bacteria such as colistin are last resort drugs. Importantly, antimicrobial peptides differ from many antibiotics in their pharmacodynamic characteristics. Here we implement these differences within a theoretical framework to predict the evolution of resistance against antimicrobial peptides and compare it to antibiotic resistance. Our analysis of resistance evolution finds that pharmacodynamic differences all combine to produce a much lower probability that resistance will evolve against antimicrobial peptides. The finding can be generalized to all drugs with pharmacodynamics similar to AMPs. Pharmacodynamic concepts are familiar to most practitioners of medical microbiology, and data can be easily obtained for any drug or drug combination. Our theoretical and conceptual framework is therefore widely applicable and can help avoid resistance evolution if implemented in antibiotic stewardship schemes or the rational choice of new drug candidates.

---

Antibiotic resistance is prevalent (1) and evolves quickly. It takes only a few years from the introduction of a new antibiotic to the clinic until resistant strains emerge(2). Prudent use and the introduction and development of novel antibiotics are currently considered to be the most effective ways to tackle resistance evolution(3). The prediction of when and how antibiotic resistance evolves and spreads is notoriously difficult, but would be extremely informative for antibiotic stewardship and the development of new drugs.

Amongst the new drugs under development are antimicrobial peptides (AMPs)(4). AMPs are peptides that have spatially explicit hydrophobic and cationic residues(5). Note that for example polymixins (including colistin) are usually subsumed under antibiotics, also fall into this category as they are AMPs of bacterial origin(6),(7). One of the alleged advantages of AMPs is that bacterial resistance would evolve much more slowly than against antibiotics(5, 8), a highly desirable property(9).

We have recently demonstrated that AMPs from multicellular organisms affect growing bacterial populations differently from antibiotics, i.e. they differ in their pharmacodynamics (or dose-response relationship)(10). A similar observation has been reported for colisitin a last resort drug to treat Pseudomonas infections(11). Pharmacodynamic characteristics of susceptible and resistant bacterial strains can be used to illustrate the selection of resistance under treatment with range of dosage(12).Such application is based on the concept of the ‘mutant selection window’ (MSW, Fig 1)(13, 14). The MSW has been successfully applied in animal models, demonstrating its value to understand resistance emergence *in vivo*(15).

**Fig 1.**
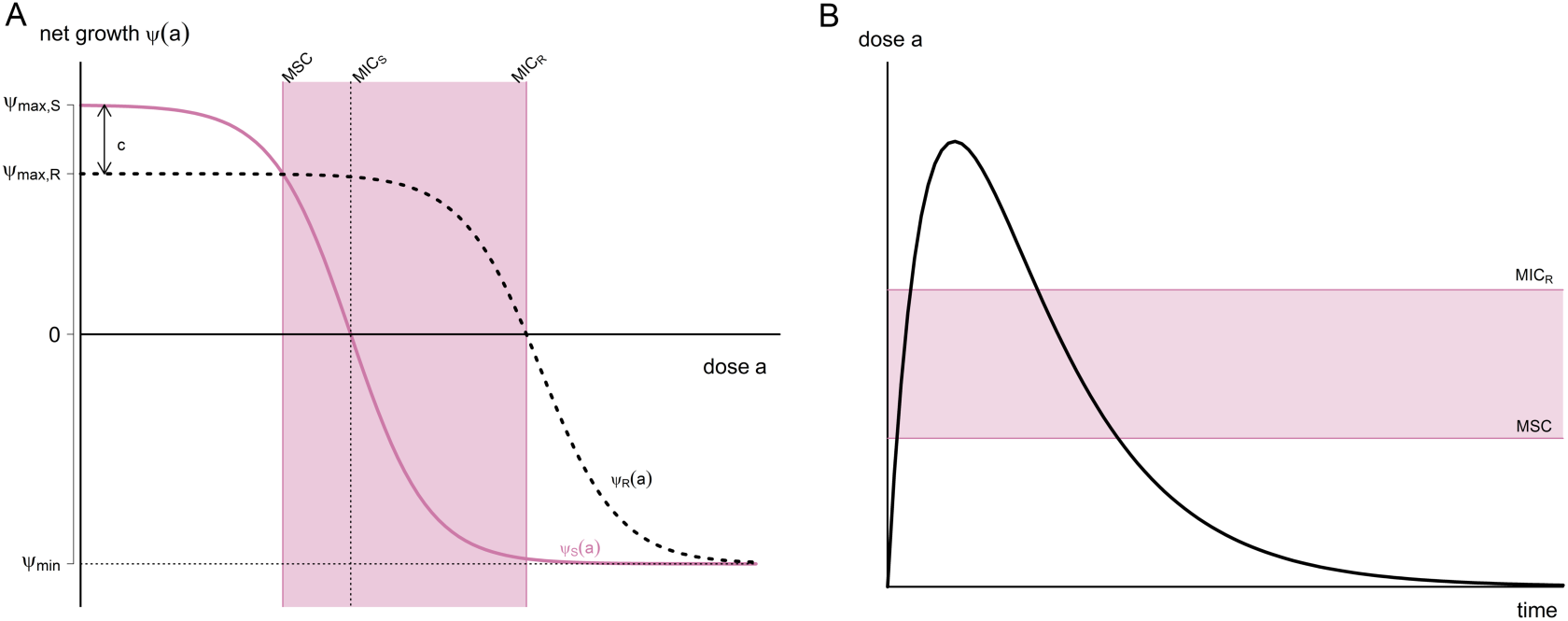
The revised mutant selection window and pharmacodynamic parameters. **(a)** The mutant selection window (MSW) is defined as the antimicrobial concentration range in which resistant mutants are selxsected (13). Following (14), we determine the MSW using net growth curves of a susceptible strain *S* and a resistant strain *R*. Mathematically, net growth is described with the pharmacodynamic function *ψ(a)* ((20), see Materials and Methods and Fig S3 for details). In short, the function consists of the four pharamcodynamic parameters: net growth in absence of antibicrobials *ψ*_*max*_, net growth in the presence of a dose of antimicrobials, which effects the growth maximal, *ψ*_*min*_, the *MIC* and the parameter *κ*, which describes the steepness of the pharamcodynamic curve. Here, the two pharmacodynamics functions *ψ*_*s*_(*a*) (continuous pink line) and *ψ*_*R*_(*a*) (dotted black line) describe the net growth of the *S* and *R*, respectively, in relation to the drug concentration *a*. Cost of resistance *c* is included as a reduction of the maximum growth rate of the resistant strain *ψ*_*max,R*_, with *c* =1 − *ψ*_*max,R*_/*ψ*_*max,s*_. Note that with this definition, cost of resistance is expressed as reduction in net growth rate in absence of antimicrobials (a = 0). The lower bound of the MSW is the concentration for which the net growth rate of the resistant strain is equal to the net growth rate of the sensitive strain and is called the minimal selective concentration (MSC) (see Materials and Methods for analytic solution, see Fig S1 for how the MSC is influenced by pharamcodynamic parameters os the sensitive strain). The upper bound is given by the MIC of the resistant strain *MIC*_*R*_.We calculate the size of the MSW as: 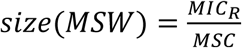. **(b)** The boundaries of the MSW applied to the pharmacokinetics of the system.

The width of the mutant selection window is partly determined by the steepness of the pharmacodynamic curve (see Fig 1). Importantly the concentration range between no killing and maximal killing is much narrower for AMPs than antibiotics, resulting in a much steeper curve. The maximum killing rate of AMPs is much higher than of antibiotics, as reflected in quicker killing time(16). Another difference relevant to the evolution of resistance is the finding that many antibiotics increase mutation rates of bacteria(17, 18),(19), but the AMPs tested so far do not show such an effect as they do not elicit bacterial DNA damage responses (17, 18).

Here we use a pharmacodynamics approach that has been widely used to describe sigmoid dose-response relationships (20, 21),(22, 23) to study the evolution of resistance of a homogeneous population. Our work uses the formulation of pharmacodynamic function from Regoes et al(20). We particularly explored how the steepness of the pharmacodynamic curve (described by the the Hill coefficient κ), together with other pharmacodynamic parameters determine the probability of resistance evolution(20). The potential importance of the Hill coefficient *κ* is often overlooked in many pharmacodynamic models, where it simply set to 1 for all drugs(24). Recent work includes the Hill coefficient (25, 26), indicating the importance of this pharmacodynamic parameter.

We use this approach with different parameter values for *κ*, derived from empirical data, as this allows us to calculate the size of the mutant selection window that generalizes over all possible resistant strains. Gullberg *et al.* demonstrated(14) that resistant mutants are already under positive selection below the MIC (minimum inhibitory concentration) of the susceptible strain. We therefore use the mutant selection concentration (MSC, Fig 1A) as the lower boundary, not the MIC of the sensitive strain that was used previously(12, 13). Using empirical parameter estimates for AMPs and antibiotics, we show that the probability of resistance evolution against AMPs (or any drug with similar pharmacodynamics properties) is much lower than for antibiotics. We therefore provide a robust and generalizable predictive framework for studying the evolution of drug resistance. This is particularly useful to apply when new drugs are introduced, i.e. before resistance has evolved.

## Results

The mutant selection window (Fig 1) shows the concentration of an antimicrobial under which susceptible strains are suppressed, but resistant strains can still grow(13). We show that the lower bound of the mutant selection window (MSC) can be calculated based solely on the pharmacodynamics of the susceptible strains and the costs of resistance (Fig 1A, Fig 2A, equation 3). The cost is defined here as the reduction of growth rate in a drug free environment.

**Fig 2.**
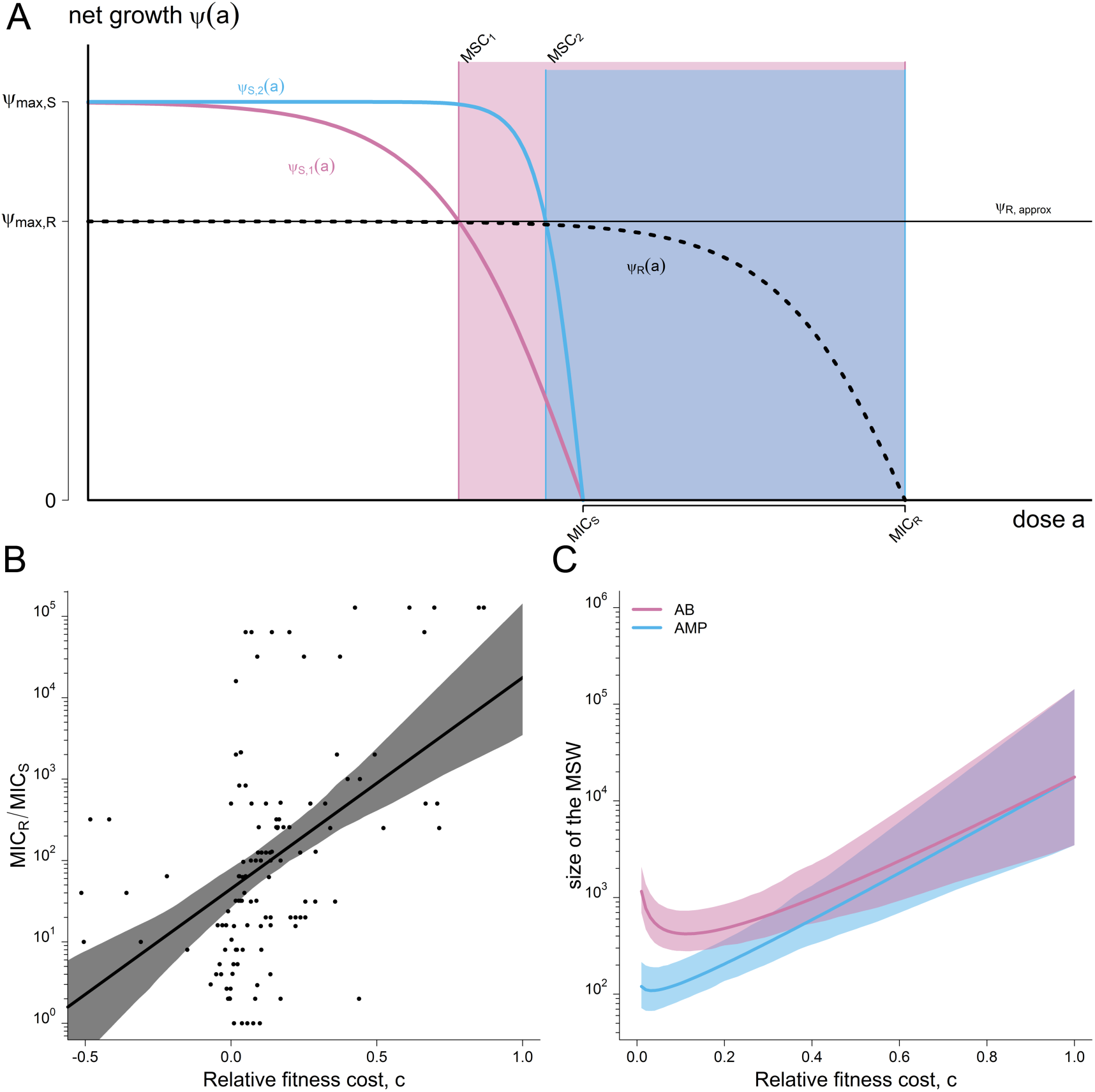
The mutant selection window for arbitrary mutant strains. The two boundaries of the MSW, MSC and MIC_R_, are influenced differently by the pharmacodynamic parameters of the sensitive strain S and the resistant strain R. **(a)** The lower boundary of the MSW (MSC) depends primarily on the pharmacodynamic parameters of the sensitive strain, assuming that the net growth rate of the resistant strain below the MSC is approximately at the same level as without antimicrobials: **ψ**_*R*_(*a*) ≈ **ψ**_*max,S*_(1 − *C*) = **ψ**_*R,approx*_, for 0 < *a* < *MSC* (**ψ**_*R*_: dotted black line; **ψ**_*R,approx*_:continuous black line) (see Materials and Methods for details). The effect of each of the four pharamcodynamic parameters and of the cost of resistance on the MSC is depicted in Fig S1. We plotted the pharmacodynamic function *ψ*_*S*_(*a*) of two sensitive strains with varying *κ* values: *ψ*_*S,1*_(*a*) representative for Abs with a small *κ* (*κ* = 1.5, pink) and *ψ*_*S,2*_(*a*) representative for AMPs with a large *κ* (*κ* = 5, blue). Increasing the *κ* value results in increasing the MSC (MSC_1_ (pink) <MSC_2_ (blue)). **(b)** The upper boundary of the MSW is per definition the *MIC*_R_, which is linked to its fitness cost, i.e. the upper boundary *MIC*_*R*_ increases with costs *c* (data from(44)). Here, the log-linear regression and the 95% confidence interval are plotted. See materials and methods for details of the statistics. **(c)** The relationship between cost of resistance, other pharmacodynamic parameters, and the size of the MSW is complex. We show that because both boundaries of the MSW - the MSC and the MIC_R_ - are influenced by costs of resistance c, the lowest MSW window size is achieved at intermediate cost of resistance *c*. We plotted the size of the MSW (line) and the 95% confidence intervals for both AMP-like and AB-like pharmacodynamics, with *ψ*_*max,S*_= 1, *MIC*_*s*_ = 1, *ψ*_*min,S,AB*_ =−5, *ψ*_*min,S,AMP*_ = −50, *κ*_*S,AB*_ = 1.5 and *κ*_*S,AMP*_ = 5. *ψ*_*max,R*_ was calculated using the relationship log_10_(MIC_R_/MIC_S_) = 2,59 * c + 1,65.

The pharmacodynamics of AMPs and antibiotics differ significantly(10): the pharmacodynamic curves of AMPs are much steeper as captured by a higher Hill coefficient *κ* (see Fig 2A); the step from a concentration with no effect to a killing concentration is therefore much smaller. This feature is likely due to a higher number of “hits” that AMPs need to deliver to bacteria to kill them and perhaps cooperative binding of AMPs molecules to the cell membrane(27). This results in a narrower MSW for AMPs than antibiotics The MSW opens at lower concentrations when the costs of resistance are low. Our re-analysis of data on antibiotic resistance against a variety of antibiotics in a number of different bacterial species (data from(28)) shows that the upper bound of the MSW correlates with the cost of resistance (Fig 2B). Taken together we are now in a position to estimate the size of the MSW for any drug, if estimates of pharmacodynamic parameters based on the sensitive strains, including the MIC, the maximum effect and the steepness of the pharmacodynamics curve are available (Fig 1A, Fig 2C).

Next we wanted to explore if the differences between AMPs and antibiotics in the width of the MSW correlated with different probabilities of drug resistance evolution within a host. A further difference between AMPs and antibiotics is that some antibiotics increase mutagenesis but AMPs do not(17, 18). We incorporated this difference in addition to the difference in the steepness of the pharmacodynamics relationship into a stochastic model describing bacterial replication and evolution under selection pressure from AMPs. We consider two cases here: (a) do resistant mutants emerge (answering this question requires a stochastic model) and (b) do resistant mutants drive the susceptible strains to extinction?

We find that resistance emerges with a much higher probability for the parameter settings of antibiotics (top row Fig 3B) than for AMPs in our simulations (bottom row Fig 3B, Fig 3A). All intermediate cases, where we simulated changes in one or two of the parameters *κ* mutation rate and maximum effect, also reduce the probability of resistance emergence compared to ‘pure’ antibiotics.

**Fig 3.**
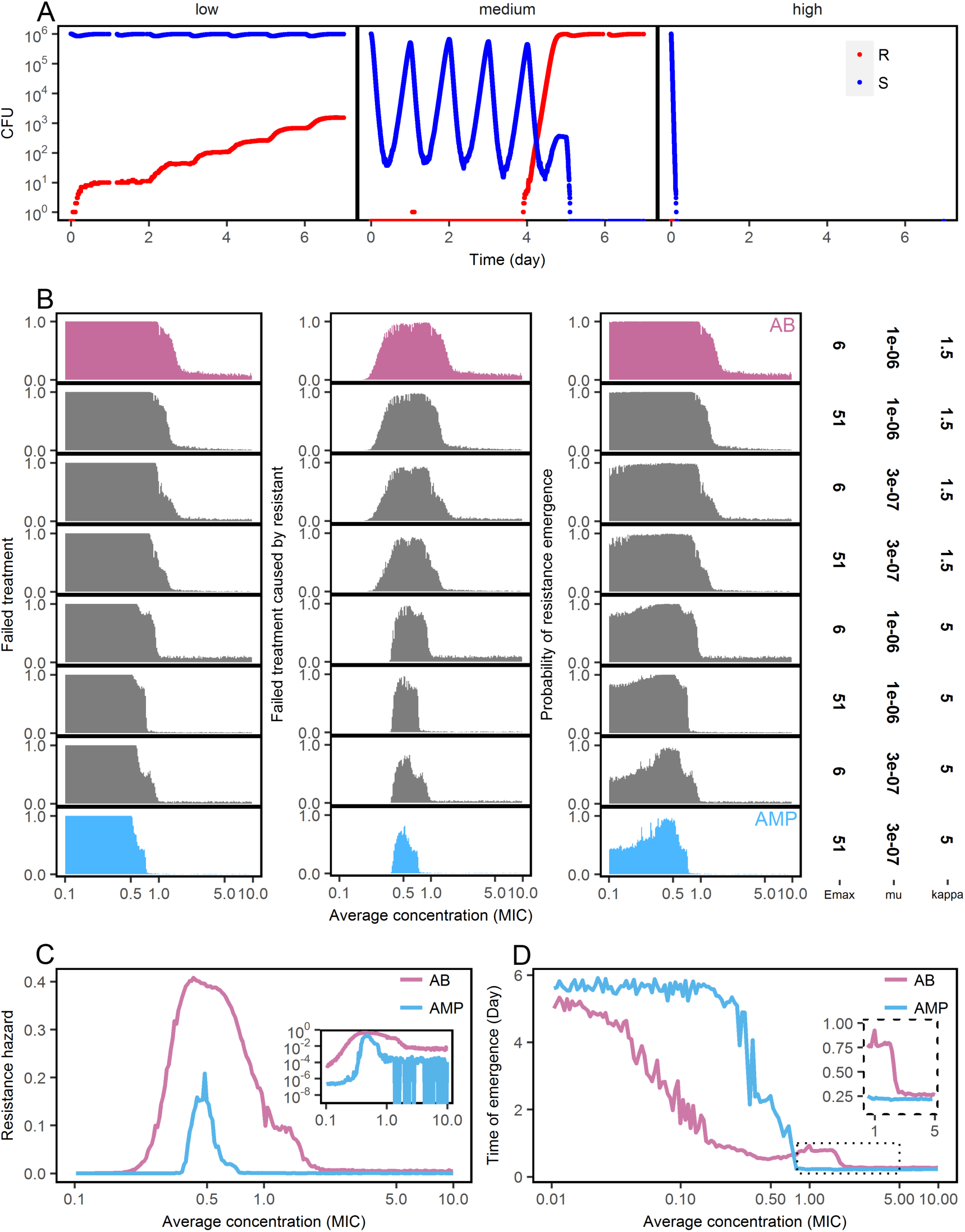
Evolution of drug resistance determined by pharmacodynamics. (a) At high dose antimicrobials achieve maximal effects and rapidly kill most of the population, preventing resistance evolution (left). At medium dose, the sensitive strain will not be eliminated immediately, and resistant mutants emerge (central). At low dose, the sensitive strain will not be removed, the mutants emerge as well, but will not quickly reach equilibrium due to substantial fitness costs (right, resistant: pink, susceptible: blue), **(b)** Simulations comparing the range from ‘pure’ antimicrobials peptides (AMP) to ‘pure’ antibiotics (AB) by altering μ, *ψ*_*min*_ and *κ*. We find that the probabilities of treatment failure (left), of failure caused by resistant strains (middle) and of resistance emergence are always higher under the AB-scenario than the AMP-scenario. A successful treatment requires less AMP than AB. **(c)** Following (29) we calculate the resistance hazard as the time-averaged proportion of mutants in a patient under a particular treatment dose. We find that AMPs are much less likely to select for resistance across concentrations than antibiotics (inset graph: a log-scale view). **(d)** Time to resistance is much longer under AMP than AB treatment when the average concentration is below MIC, but shorter around MIC and equal in higher concentrations (inset graph). The parameters are: *ψ*_*max,S*_ = 1, *ψ*_max,R_ = 0.9, *κ*_*AB*_ = 1.5, *κ*_*AMP*_ = 5, *ψ*_*min,AB*_ = −5, *ψ*_*min,AMP*_ = −50, *MIC*_*S*_ = 10, *MIC*_*R*_ = *MIC*_*S*_ * 10^[2.59 *(*μ*_*max,S*_-*ψ*_*max,R*_)+ 1.65]^. *μ*_*AB*_ = 10^−6^, *μ*_*AMP*_ = 3 * 10^−7^, *k*_*a*_ = 0.5, *k*_*e*_ = 0.2, *d*_*n*_ = 0.01, *τ* = 1/24.

We also find that resistant mutants are much more likely to drive the susceptible bacterial populations to extinction under antibiotic than under AMP treatment (Fig 3 B). Again, this result also holds when we study intermediate cases. In summary, our results show that the application of drugs with low *κ*, mutation elevation and low maximum effect, i.e. characteristics found in most common antibiotics, inherently bears a high risk of causing the evolution of resistance.

We have shown before(10) that combinations of AMPs have higher *κ* and lower MICs than individual AMPs. This also results in differences in resistance selection and the extinction of susceptible strains, consistent with the results above.

Day *et al* (29) provided an approach to calculate a resistance hazard: a measure that combines the time of resistance emergence and its selection within a host. We calculated similar resistance hazard for AMPs in comparison to antibiotics. The simulation results show (Fig 3C) that the hazard is much higher and the concentration range much wider under antibiotic treatment than under AMP treatment. Also, when resistance evolves, it emerges earlier in the antibiotic scenario than in the AMP scenario at low concentrations (Fig 3D). In certain concentrations (for example, around MIC in our simulation), resistance emerges earlier in AMP than in antibiotics (Fig 3D). Time of emergence is mostly affected by *κ* and mutation rate: higher *κ* and lower mutation rate, the latter more important when population sizes are small, confer delayed resistance emergency (Fig S4).

## Discussion

Our predictions suggest that AMPs, or in fact any antimicrobial drug with similar pharmacodynamics, are much less likely to select drug-resistant mutants than antimicrobials with antibiotic-like characteristics. Our theory is blind to the molecular mechanism of action but captures the dynamically relevant aspects of action.

We assume that pharmacodynamics and mutagenic properties of AMPs are significantly different from antibiotics. This assumption is based on limited data of AMPs in the literature(10, 17). More experiments with a variety of antimicrobial peptides are needed to determine if AMP like characteristics can be indeed generalized and if these characteristics are significant different from antibiotics.

In the light of our results, increasing *κ* and/or the maximum effect are desirable for any drug as well as advantageous to hosts managing their microbiota using AMPs. Our model therefore provides useful information for the development of new antimicrobial drugs: higher *κ* and maximum effect will impose much weaker selection on the bacteria to evolve resistance in lower concentrations, and clear the bacterial population more quickly in higher concentration which will, in turn, reduce the probability of resistance emergence. Currently mostly AMPs display these properties, but it is likely that new antibiotics that target the cell membrane or wall display similar pharmacodynamics.

The smaller MSW under AMPs is a direct consequence of the steeper pharmacodynamic functions(10). It is important to note that this relationship hinges on the realization that the window opens at the concentration at which the resistant strains have a higher growth rate than the sensitive strain, well below the MIC of the sensitive strain(14). Thus, a high Hill coefficient (*κ*) would constitute a promising characteristic of new antimicrobials. The other characteristics in which AMPs differ from antibiotics - mutagenesis and maximum effect - affect mostly the time until resistance emerges, but not the size of the MSW. Because this time becomes shorter with higher population sizes, these characteristics may have less significance for clinical infections (30).

We find that time to resistance emergence in AMPs is longer than in antibiotics when the concentration is low (subMIC). Around MIC resistance against AMPs seems to emerge quicker than against antibiotics (FIG D). This counterintuitive result is explained by the fast removal of the sensitive strains caused by the combination of high *κ* and low psimin and is not related to the mutation rate *per se*. Overall the probability of resistance emergence is lower for AMPs as higher concentrations quickly remove the sensitive population. Chevereau *et al*. (31) reached a different conclusion using a different modeling approach. They modeled the pharmacodynamics only for positive growth and continuously adjusted the drug concentration to maintain the overall growth rate at half of the maximal in the simulation. In this scenario, drugs with sensitive dose-response would facilitate evolution due to the wide distribution of fitness, a scenario that seems unlikely in real antimicrobial treatment.

One recommendation derived from our modeling approach is that drugs that show pharmacodynamics resembling AMPs should be good candidates for slowing the evolution of resistance. Interestingly, combinations of AMPs result in increased κ, which our model predicts to bear lower risks of evolution of resistance(10). It is often argued that combination therapy reduces resistance evolution (but also see (32)), as it is supposedly more difficult to evolve resistance against more than one mechanism at a time. Our approach indicates that combination therapy might even prove effective if there are mutations that confer complete cross-resistance to the drugs in the combination.

It has been proposed that bacterial resistance evolution against AMPs is highly unlikely (5, 8). Yet, *in vitro* experimental evolution has demonstrated that resistance to AMPs can arise (33–35) and AMP-resistance mechanisms have been characterized (36). Against antibiotics, resistance can increase the MIC by 2-3 orders of magnitude in a relatively small bacterial population(37), a range that has not been observed for AMPs. Though AMPs provide promising leads for drug development (4), their conserved killing mechanisms also argue for caution. In their paper ‘arming the enemy’, Bell et al.(38) discussed the high likelihood of cross-resistance against, for example, human AMPs. This problem has hardly been studied. Our analysis suggests how one could reap the benefits of AMPs without arming the enemy: we should rely on agents with AMP-like pharmacodynamics. This in principle can be adopted without using AMPs themselves.

Pharmacodynamic estimates can be easily and routinely obtained from time-kill curves. This can also be achieved for drug combinations(10). A report by the *Leopoldina*, the German National Academy of Sciences, recently recommended to use new drugs only in combination to avoid fast resistance evolution(39). The scientific support for this notion is limited and controversial(32, 40, 41). In clinical situations pharmacodynamic approaches can provide a first informed guess. Also, the risk of resistance evolution based on the pharmacodynamics of drug candidates will be a useful additional criterion to develop new drugs. We would also like to note that the concept of the mutant selection window has been applied to understand antiviral resistance evolution(42), and hence our approach has the potential to inform antiviral resistance research and ultimately treatment as well.

In order to generate predictions on resistance evolution based on pharmacodynamics, one of our main goals of the project, we made a number of simplifying assumptions. The pharmacodynamics are based on data of initial killing only. Moreover, we assume homogeneous populations over time and space. Expanding the framework to integrate tolerance and resistance is possible but would require pharmacodynamic estimates and additional functions. Another possible extension of our work would be to include pharmacodynamic estimates of resistant strains that change over time due to compensatory mutations and to cross resistance or collateral sensitivity when exposed to combinations of antimicrobials. Finally, we assumed the same pharmacokinetics for all cases in our study. As AMPs are currently rarely used (Colistin being the notable exception), future empirical work will inform realistic parameter estimates for pharmacokinetics. In all cases however, the basis of any analysis concerning resistance evolution is the influence of individual pharmacodynamic parameters, for which we provide a framework.

## Materials and Methods

For the parameterization of the predictive models, we used two main sources. The pharmacodynamic parameters are taken from one of our own studies that determines pharmacodynamics for AMPs and antibiotics under standardized conditions(10). In short, time kill experiments with different AMP concentrations were conducted and the slopes of the linear regressions were used to calculate the parameters of the pharmacodynamic function. Here, we only took into account the intial kill rates and assumed a homogeneous population structure. The estimates of mutation rates again are from our own comparative study on mutagenesis under AMP and AB treatment(17).

### Calculation of the size of the mutant selection window

The size of the mutant selection window (MSW) depends on the lower and upper bound of the MSW and is calculated as

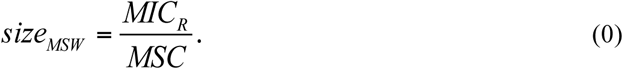

The lower bound of the MSW is the concentration for which the net growth rate of the resistant strain is equal to the net growth rate sensitive strain and is called the minimal selective concentration (MSC). The upper bound of the MSW is the MIC of the resistant strain (MIC_R_) (Fig 1 A).

To analytically describe the MSW, we use the pharmacodynamic (PD) function *ψ*(*a*), which mathematical describes the net growth rate with a Hill function:

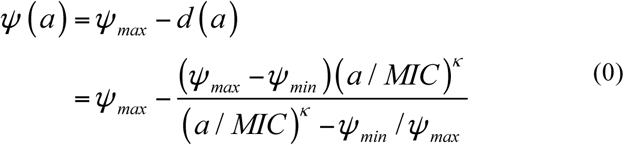

((10, 20, 21)). Here, *a* is the antimicrobial drug concentration, *ψ*(*a* = 0) = *ψ*_*max*_, *d*(*a*) is the effect of the antimicrobial with the dose *a*, and *ψ*(*a* → ∞) = *ψ*_*min*_. Therefore, the maximal effect *E*_*max*_ is *E*_*max*_ = *ψ*_*max*_ - *ψ*_*min*_. The parameter *MIC* denotes the concentration that results in zero net growth (this definition differs from the "official" MIC definition by Mouton et al (43)). The Hill coefficient *κ* describes the steepness of the curve; functions with higher *κ* describe steeper curves (Fig 2A). For illustration of the pharamcodynamic parameters see Fig S3). Cost of resistance *c* is included as a reduction of the maximum growth rate of the resistant strain in absence of antimicrobials with *c* = 1-*ψ*_*max,R*_/*ψ*_*max,S*_ (Fig 1A, 2A). The pharamcodynamic function can be described for both a drug susceptible strain *S* and a drug-resistant strain *R*, with *ψ*_*S*_(*a*) and *ψ*_*R*_(*a*), respectively. The *MSC* is calculated as *ψ*_*S*_(*a*) = *ψ*_*R*_(*a*). We assume that the net growth rate of the resistant strain below the *MSC* is, for any given concentration a, with 0 < *a* < *MSC*, approximately at the same level as without antimicrobials and therefore we set *ψ*_*R*_(*a*) ≈ *ψ*_*R*,*approx*_ (illustrated in Fig 2A). With *ψ*_*R,approx*_. = *ψ*_*max,R*_ = *ψ*_*maxS*_ (1-*c*), we are able to describe the net growth rate of the resistant strain with the net growth rate of the sensitive strain *ψ*_*max,S*_ and the costs of resistance c: *ψ*_*R*_(*a*) ≈ *ψ*_*R,approx*_ = *ψ*_*max,S*_ (1-*c*). This is valid because *MIC*_*R*_≫ *MIC*_*S*_ and assuming *κ*_*R*_ >≈ *κ*_*S*_. The analytic solution of the *MSC* is

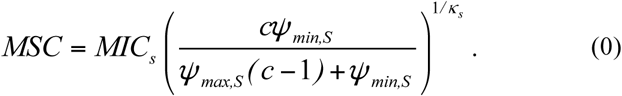

### Analysis of the relationship between cost of resistance c and MIC_*R*_

Data(44) determining relationship between fitness of resistant strains and *MIC*_*R*_/*MIC*_*C*_ swas re-analyzed. The dataset contained information about increase of MIC due to resistance and fitness of the resistant strain. The dataset summarizes cases of bacterial resistance to antibiotics. Similar data for AMPs have been compiled recently(30) but are yet too scarce to include in the following analysis. We therefore assumed similar relationships for both antibiotics and AMPs. We calculated cost of resistance *c* as c = 1 - fitness, using n= 128 observations compiled in the mentioned dataset. Fitting a log_10_ transformed linear regression to the data resulted in the parameterized function log_10_(MIC_*R*_/*MIC*_*S*_) = 2,59 * c + 1,65, (R^2^ = 0.22). The data was then resampled with using bootstrapping to (i) determine the 95% confidence interval of log-linear regression of the data as interval, where 95 % of the regression fall into (see fig. 2B) and (ii) to include the variance of the data when determining the size of the mutant selection window (MSW)(see fig. 2C). For the latter, the given dataset was fitted to the mentioned log-linear regression 200 times, resulting in 200 parameter sets for the regression. Each parameter set was then used to calculate the size of the MSW depending on the cost of resistance. The 95% confidence interval was then calculated as the interval, in which 95% of the calculated size of the MSW are in for a given cost.

### Model of evolution and prediction of resistance

To study resistance evolution we used a mathematical model that incorporates pharmacodynamics (PD) and pharmacokinetics (PK) and captures population dynamics of bacterial populations under treatment with antimicrobial drugs(20). We ran stochastic simulations to calculate the probability of resistance emergence, the probability of the resistant strain, the time to resistance emergence and the risk of resistance (the resistance hazard(29)).

To simulate treatment, we consider a patient harboring 10^6^ susceptible bacteria. Bacterial mutation rates are assumed to depend on the antimicrobial used for treatment (antibiotics or AMPs). When a resistant strain arises it is assumed to have an MIC ten-fold that of susceptible wild-type strain. For simplicity, we only consider one type of mutant. Antimicrobials are administered every day (see Supplement for pharmacokinetics), and treatment lasts one week.

The population dynamics of the susceptible and resistant strains is captured in the following system of differential equations:

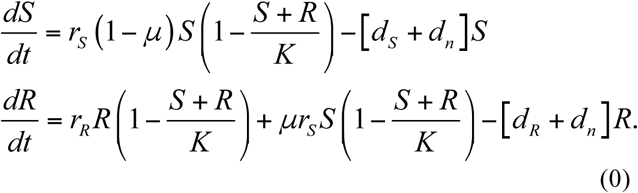

Where *S* represents the wild-type strain and *R* represents the resistant strain. The maximum net growth rate *ψ*_*max*_ is the difference between the replication rate *r* and the intrinsic death rate *d*_*n*_: *ψ*_*max*_ = r-d_n_. *μ* is the mutation rate.

To include the change of antimicrobial concentrations over time (pharmacokinetics) into our mode, we define the death rate to be dependent on the time-dependent antimicrobial concentration a(t):

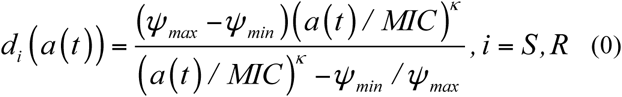

We assume a time-dependent pharmacokinetic function *a(t)* of the following form (see also Fig S2):

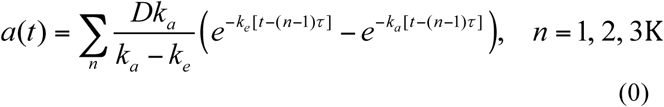

Here, *k*_*a*_ *is* the absorption rate, and *k*_*e*_ is the decay rate. *D* is the dose given each time, *n* is the number of doses, *τ* is the dosing frequency. We define the treatment dose as the average concentration in the course of treatment:

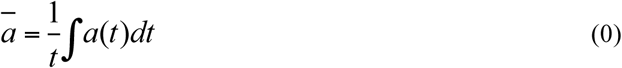

We implemented the model in Equation 4 stochastically using the Gillespie algorithm(45), which allowed us to monitor how frequently mutants arise. Parameters were selected based on empirical data as stated above. The net growth rate of wild-type in the absence of antimicrobials was set as 1. Mutants suffer fixed or resistant-level related costs (see Fig 2). *κ* of AMPs and antibiotics were set as 5 and 1.5, respectively (10). *ψ*_min_ for AMPs is fixed as −50 hour^-1^; and for antibiotics is fixed as - 5 hour^-1^. Mutation rates in AMPs are assumed to be three times lower than in antibiotics, in accordance with our empirical estimates (17). All the parameters and their values are listed in Table S1. All the pharmacokinetic parameters are the same in different simulations (see Fig S2). For each set of parameters, cohorts of five hundred infected individuals were simulated. Successful treatment is defined as complete clearance of both sensitive and resistant strains at the end of the one-week treatment. For each cohort, we calculate the probability of treatment success as the proportion of individuals in whom treatment was successful. In each individual, we score the time of emergence of resistance strains, and estimate the resistance hazard based on the average probability of treatment success and the population size of bacteria over time. The hazard function can be written as,

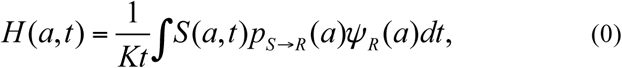

where *K* is the capacity, *S* denotes population size of sensitive strain and *p*_*S→R*_ is probability of a treatment developing resistance, which is calculated from the results of simulations, *ψ*_*R*_ is the growth rate of resistant strain. Our hazard function calculates the average proportion of resistant population under certain treatment dose and duration.

### Implementation

The analysis was performed in R (v. 3.1.3&v. 3.2.2) (46) using RSTUDIO (v. 0.98.1103&0.99.903) ^35^. The code is available upon request.

## Acknowledgements

We are grateful to Olivia Judson and Sebastian Bonhoeffer for comments on the manuscript.

## Author contributions

All authors participated in the design and interpretation of the results. GY was primarily responsible for the predictive modelling, DYB for the PDwork. All authors contributed to the writing of the paper. JR wrote the first draft, RRR led the mathematical work.

## Data and materials availability

The model will be made available as a remarkup document for use.

## References

1. Laxminarayan R, Sridhar D, Blaser M, Wang M, Woolhouse M (2016) Achieving global targets for antimicrobial resistance. Science (80-) 353: 874–875.

2. McClure NS, Day T (2014) A theoretical examination of the relative importance of evolution management and drug development for managing resistance. Proc Biol Sci 281: 20141861.

3. World Health Organization (2014) The evolving threat of antimicrobial resistance: Options for action. WHO Publ:1–119.

4. Czaplewski L et al. (2016) Alternatives to antibiotics — a pipeline portfolio review. Lancet Infect Dis 16:239–251.

5. Zasloff M (2002) Antimicrobial peptides of multicellular organisms. Nature 415:389–395.

6. Hancock REW, Sahl H-G (2006) Antimicrobial and host-defense peptides as new anti-infective therapeutic strategies. Nat Biotechnol 24:1551–7.

7. Jochumsen N et al. (2016) The evolution of antimicrobial peptide resistance in Pseudomonas aeruginosa is shaped by strong epistatic interactions. Nat Commun 7:13002.

8. Fjell CD, Hiss JA, Hancock REW, Schneider G (2012) Designing antimicrobial peptides: form follows function. Nat Rev Drug Discov 11:37–51.

9. Ling LL et al. (2015) A new antibiotic kills pathogens without detectable resistance. Nature. 517:455–459

10. Yu G, Baeder DY, Regoes RR, Rolff J (2016) Combination Effects of Antimicrobial Peptides. Antimicrob Agents Chemother 60:AAC.02434-15.

11. Mohamed AF, Cars O, Friberg LE (2014) A pharmacokinetic/pharmacodynamic model developed for the effect of colistin on Pseudomonas aeruginosa in vitro with evaluation of population pharmacokinetic variability on simulated bacterial killing. J Antimicrob Chemother 69:1350–1361.

12. Firsov AA et al. (2013) Bacterial resistance studies using in vitro dynamic models: The predictive power of the mutant prevention and minimum inhibitory antibiotic concentrations. Antimicrob Agents Chemother 57:4956–4962.

13. Drlica K, Zhao X (2007) Mutant selection window hypothesis updated. Clin Infect Dis 44:681–8.

14. Gullberg E et al. (2011) Selection of Resistant Bacteria at Very Low Antibiotic Concentrations. PLoS Pathog 7:e1002158.

15. Cui J et al. (2006) The mutant selection window ub Rabbits Infected with Staphylococcus aureus. J Infect Dis 194:1601–1608.

16. Fantner GE, Barbero RJ, Gray DS, Belcher AM (2010) Kinetics of antimicrobial peptide activity measured on individual bacterial cells using high-speed atomic force microscopy. Nat Nanotechnol 5:280–5.

17. Rodríguez-Rojas A, Makarova O, Rolff J (2014) Antimicrobials, Stress and Mutagenesis. PLoS Pathog 10:e1004445.

18. Rodríguez-Rojas A, Makarova O, Müller U, Rolff J (2015) Cationic Peptides Facilitate Iron-induced Mutagenesis in Bacteria. PLOS Genet 11:e1005546.

19. Kohanski M, DePristo M, Collins JJ (2010) Sublethal antibiotic treatment leads to multidrug resistance via radical-induced mutagenesis. Mol Cell 37:311–20.

20. Regoes RR et al. (2004) Pharmacodynamic Functions: a Multiparameter Approach to the Design of Antibiotic Treatment Regimens. Antimicrob Agents Chemother 48:3670–3676.

21. Shen L et al. (2008) Dose-response curve slope sets class-specific limits on inhibitory potential of anti-HIV drugs. Nat Med 14:762–766.

22. Bonapace CR, Friedrich L V., Bosso JA, White RL (2002) Determination of antibiotic effect in an in vitro pharmacodynamic model: Comparison with an established animal model of infection. Antimicrob Agents Chemother 46:3574–3579.

23. Corvaisier S et al. (1998) Comparisons between antimicrobial pharmacodynamic indices and bacterial killing as described by using the Zhi model. Antimicrob Agents Chemother 42:1731–1737.

24. Craig WA (1998) Pharmacokinetic / Pharmacodynamic Parameters: Rationale for Antibacterial Dosing of Mice and Men Author. Clin Infect Dis 26:1–10.

25. Nielsen EI, Friberg LE (2013) Pharmacokinetic-Pharmacodynamic Modeling of Antibacterial Drugs. Pharmacol Rev:1053–1090.

26. Sy SKB, Derendorf H (2014) in Applied Pharmacometrics, eds S S, Derendorf H (American Association of Pharmaceutical Scientists), pp 229–257.

27. AV H (1910) The possible effects of the aggregation of the molecules of hæmoglobin on its dissociation curves. J Physiol 40:4–7.

28. Melnyk AH, Wong A, Kassen R (2014) The fitness costs of antibiotic resistance mutations. Evol Appl 8:273–283

29. Day T, Read AF (2016) Does High-Dose Antimicrobial Chemotherapy Prevent the Evolution of Resistance? PLOS Comput Biol 12:e1004689.

30. Andersson DI, Hughes D, Kubicek-Sutherland JZ (2016) Mechanisms and consequences of bacterial resistance to antimicrobial peptides. Drug Resist Updat 26:43–57.

31. Chevereau G et al. (2015) Quantifying the Determinants of Evolutionary Dynamics Leading to Drug Resistance. PLOS Biol 13:e1002299. A

32. Pena-Miller R et al. (2013) When the Most Potent Combination of Antibiotics Selects for the Greatest Bacterial Load: The Smile-Frown Transition. PLoS Biol 11:e1001540.

33. Perron GG, Zasloff M, Bell G (2006) Experimental evolution of resistance to an antimicrobial peptide. Proc Biol Sci 273:251–6.

34. Habets MGJL, Brockhurst M a(2012) Therapeutic antimicrobial peptides may compromise natural immunity. Biol Lett 8:416–8.

35. Dobson AJ, Purves J, Kamysz W, Rolff J (2013) Comparing Selection on S. aureus between Antimicrobial Peptides and Common Antibiotics. PLoS One 8:e76521.

36. Joo H-S, Fu C, Otto M (2016) Bacterial Strategies of Resistance to Antimicrobial Peptides. Phil Trans R Soc B 371:20150291

37. Barbosa C et al. (2017) Alternative evolutionary paths to bacterial antibiotic resistance cause distinct collateral effects. Mol Biol Evol:1–16.

38. Bell G (2003) Arming the enemy: the evolution of resistance to self-proteins. Microbiology 149:1367–1375.

39. Akademie der Wissenschaften Hamburg (2013) Antibiotika-Forschung: Probleme und Perspektiven (Walter de Gruyter).

40. Imamovic L, Sommer MO (2013) Use of collateral sensitivity networks to design drug cycling protocols that avoid resistance development. Sci Transl Med 5:204ra132.

41. Holmes AH et al. (2016) Understanding the mechanisms and drivers of antimicrobial resistance. Lancet 387:176–187.

42. Rosenbloom DIS, Hill AL, Rabi SA, Siliciano RF, Nowak M (2012) Antiretroviral dynamics determines HIV evolution and predicts therapy outcome. Nat Med 18:1378–1385.

43. Mouton JW, Dudley MN, Cars O, Derendorf H, Drusano GL (2005) Standardization of pharmacokinetic/pharmacodynamic (PK/PD) terminology for anti-infective drugs: An update. J Antimicrob Chemother 55:601–607.

44. Melnyk AH, Wong A, Kassen R (2015) The fitness costs of antibiotic resistance mutations. Evol Appl 8:273–283.

45. Pineda-Krch M (2008) GillespieSSA: Implementing the Stochastic Simulation Algorithm in R. J Stat Softw 25:1–18.

46. R. a language for statistical computing. Vienna 2015, A. R. F. for S. C. R: a language and for statistical computing. (http://www.Rproject.org).

